# Monomer and dimer structures of cytochrome *bo*_*3*_ ubiquinol oxidase from *Escherichia coli*

**DOI:** 10.1101/2022.12.09.519786

**Authors:** Yirui Guo, Elina Karimullina, Tabitha Emde, Zbyszek Otwinowski, Dominika Borek, Alexei Savchenko

## Abstract

The *E. coli* cytochrome *bo*_*3*_ ubiquinol oxidase is a four-subunit heme-copper oxidase that serves as a proton pump in the *E. coli* aerobic respiratory chain. Despite many mechanistic studies on this protein, it is unclear whether this ubiquinol oxidase functions as a monomer, or as a dimer in a manner similar to its eukaryotic counterparts – the mitochondrial electron transport complexes. In this study, we determined the monomeric and dimeric structures of the *E. coli* cytochrome *bo*_*3*_ ubiquinol oxidase reconstituted in amphipol by cryogenic electron microscopy single particle reconstruction (cryo-EM SPR) to a resolution of 3.15 Å and 3.46 Å, respectively. We have discovered that the protein can form a dimer in C2 symmetry, with the dimerization interface maintained by interactions between the subunit II of one monomer and the subunit IV of the other monomer. Moreover, the dimerization does not induce significant structural changes in each monomer, except the movement of a loop in subunit IV (residues 67–74).

## Introduction

The *E. coli* cytochrome *bo*_*3*_ ubiquinol oxidase is a heme-copper oxidase that reduces molecular oxygen to water while pumping protons across the membrane (**Figure 1**) (1-3). The protein has four subunits and catalyzes the four-electron reduction of O_2_ to H_2_O. The natural substrate of this enzyme is ubquinol-8. The redox center of the oxidase is located in subunit I, as shown in **Figure 1B**, with all the cofactors represented in their exact positions in the solved structure. During catalysis, electrons move in the direction from ubiquinol to heme *b* to heme *o*_*3*_/Cu ion (Cu_B_). Protons produced in the reaction are released to the positive side (P-side) of the membrane and are subsequently converted to energy through the aerobic respiratory chain (4).

**Figure 1.**
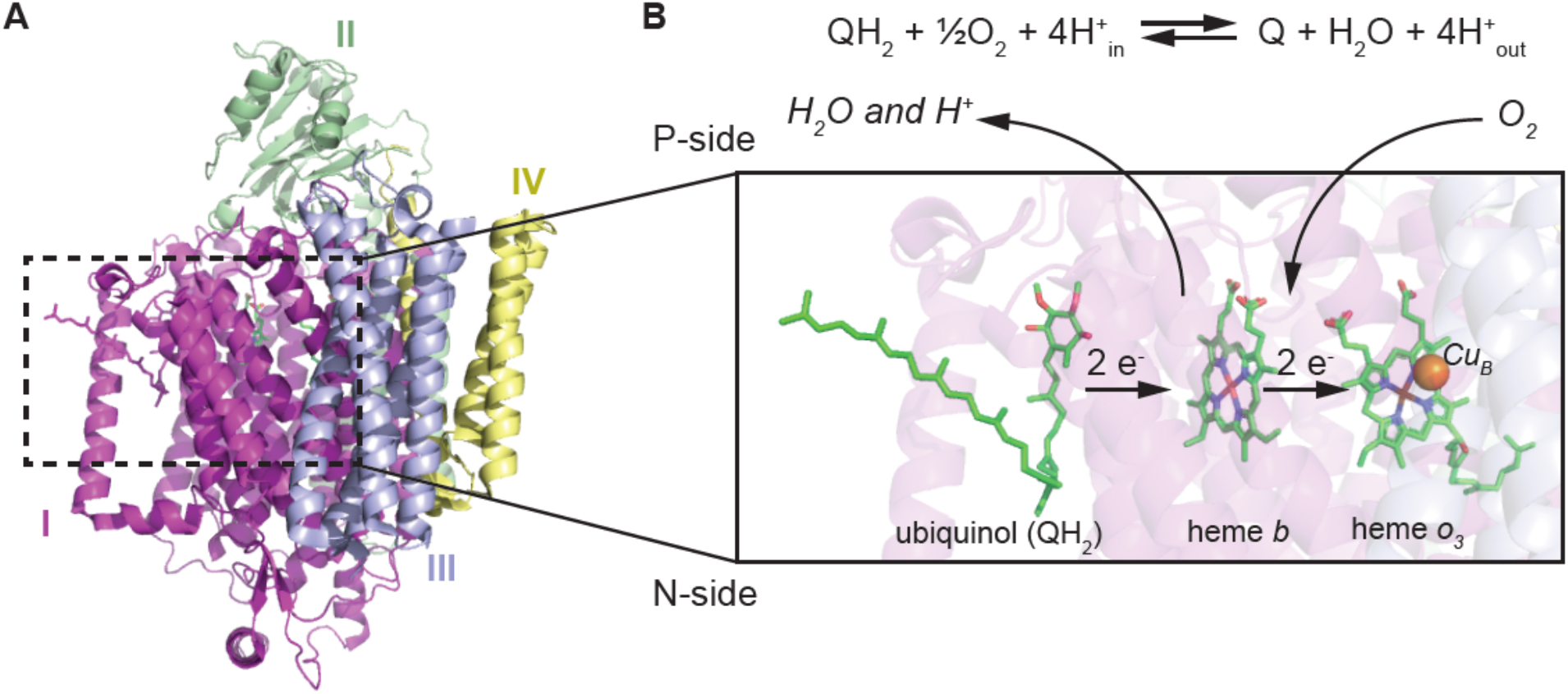
Structural and functional overview of the *E. coli* ubiquinol oxidase (adapted from PDB 1FFT and 6WTI) **A)** Structure of the *E. coli* ubiquinol oxidase with the four subunits (I, II, III and IV) shown in different colors. Black dashed box indicates the redox center of the protein. **B)** Schematic representation of electron/proton transfer at the redox center. Briefly, all redox centers (ubiquinol, heme *b*, heme *o*_*3*_ and a copper ion) are bound to subunit I, the largest subunit of the oxidase. Oxidation of one ubiquinol releases two protons on the positive side (P-side) of the membrane (2H^+^ chemistry). This reaction also creates a proton motive force which pumps two additional protons across the membrane (2H^+^ translocation).

The *E. coli* cytochrome *bo*_*3*_ ubiquinol oxidase is often compared to its eukaryotic counterparts, the mitochondrial electron transport complexes such as the mitochondrial cytochrome *bc*_*1*_ and cytochrome *c* oxidase. Both mitochondrial electron transport complexes crystalized as dimers and the dimer is believed to be important for proton translocation (5-8). The dimerization of the mitochondrial electron transport complexes is also sensitive to detergent solubilization. For example, cytochrome *c* can switch from a mixture of monomers and dimers to monomers only as detergent concentration increases (9).

However, the oligomerization status of the *E. coli* cytochrome *bo*_*3*_ ubiquinol oxidase is less clear. To date, all published structures of this ubiquinol oxidase are monomeric with only a few structural differences between each other (1-3). Firstly, the structure solved by X-ray crystallography is missing density in subunit I (residues 552-656) compared to the structures solved by Cryo-EM (1). Secondly, the crystal structure lacks ubiquinone molecule bound to the protein due to the detergent used in the crystallization conditions, while both cryo-EM structures show one ubiquinone molecule bound to subunit I (10). Several biochemical and biophysical characterizations of the enzyme resulted in contradictory observations of the oligomerization states. Musatov *et al*. (11) measured the oligomerization status by sedimentation velocity and equilibrium experiments and reported that the detergent-solubilized *E. coli* cytochrome *bo*_*3*_ ubiquinol oxidase is either monomeric or is highly aggregated; while another study by Stenberg *et al*. (12) reported the presence of ubiquinol oxidase dimer detected by blue native polyacrylamide gel electrophoresis (BN-PAGE). The dimer measured by BN-PAGE is roughly 50-fold less abundant than the monomer and is likely sensitive to detergent solubilization.

With the recent developments in cryo-EM SPR which facilitate heterogeneous sample processing and AI-based particle picking (13, 14), we were able to isolate and computationally enrich the rare dimeric ubiquinol oxidase particles from the sample that was dominated by the highly abundant monomeric particles and solved both structures, including the first dimeric structure of this protein.

## Methods

### Protein expression and purification

The *E. coli* cytochrome *bo*_*3*_ ubiquinol oxidase in this study is a byproduct (impurity) of the expression and purification of a recombinant *E. coli* ZapG/RodZ complex. The N-terminally 6His-TEVcleavage site-tagged *E. coli* ZapG (MCS1) and untagged RodZ (MCS2) genes in the petDUET1 vector were transformed into C43(DE3) competent cells. Transformants were used to inoculate 3 L of LB media with ampicillin (100 μg/ml) at 37 °C until OD600 reached 0.6. The culture was then incubated on ice for 1 h, induced with 1 mM isopropyl-β-D-thiogalactopyranoside (IPTG) and grown overnight at 20 °C. Cells were harvested by centrifugation, resuspended with 1× PBS pH 7.4 and lysed using serial freezing and thawing cycles in the presence of phenylmethylsulfonyl fluoride (PMSF, 0.5 mM), DNase1 (20 μg/ml), lysozyme (1 mg/ml), and TCEP (0.5 mM). All further purification steps were conducted at 4 °C. Cells were processed in a cell disruptor (Avastin) at 15,000 psi, 3 passes. Cell debris was spun down at 13,500×g for 20 min. The supernatant was further ultracentrifuged for 1 h 40 min at 40,000 rpm. The pellet (crude membrane preparation) was washed with 1× PBS containing 0.5 mM TCEP and ultracentrifuged again at 40,000 rpm for 1 h 30 min. Isolated membrane pellet was frozen and kept at -80 °C until the next step. The crude membrane preparation was resuspended in the extraction buffer (50 mM Na Phosphate pH 7.6, 300 mM NaCl, 20% (v/v) glycerol, 0.5 mM TCEP, 1% DDM) and incubated for 2 h. After ultracentrifugation (40 min 40,000 rpm), the supernatant was incubated overnight with 3 ml Ni-NTA beads (Qiagen) preequilibrated with the extraction buffer. The next day, the beads were washed with 30 ml of wash 1 (50 mM Na Phosphate pH 7.6, 500 mM NaCl, 15% (v/v) glycerol, 0.5 mM TCEP, 40 mM Imidazole, 0.1% (w/v) DDM) and wash 2 (50 mM Na Phosphate pH 7.6, 300 mM NaCl, 10% (v/v) glycerol, 0.5 mM TCEP, 50 mM imidazole, 0.05% (w/v) DDM) buffers and then eluted with 15 ml of the elution buffer (50 mM Na Phosphate pH 7.6, 300 mM NaCl, 5% (v/v) glycerol, 0.5 mM TCEP, 0.05% (w/v) DDM, 300 mM imidazole). The eluate was dialyzed overnight with TEV protease (purified in-house) in dialysis buffer (50 mM Tris pH 8.0, 300 mM NaCl, 5% (v/v) glycerol, 0.5 mM TCEP, 0.05% (w/v) DDM). The next day, an immobilized metal affinity chromatography (IMAC) column packed with Ni-NTA beads (Qiagen) was used for the second time where the flow-through fraction was passed 5 times through the column so that all His-tag labeled molecules were retained. Amphipol A8-35 (Anatrace) was added to the flow-through in a 1:3 (w/w) protein:amphipol ratio and incubated with gentle rotation. After 4 hours of rotation, Bio-Beads (Biorad) (15 mg beads per 1 ml of protein solution) were added and incubated overnight. The next day, the sample with proteins stabilized in amphipol was poured over an empty column to remove the Bio-Beads and concentrated using a 100 kDa cut-off centrifugal concentration device (Amicon). The concentrated sample was loaded on a Superdex 200 Increase 10/300 column equilibrated with detergent free buffer (50 mM Tris pH 8.0, 150 mM NaCl, 0.5 mM TCEP). The peaks’ fractions were collected, concentrated, aliquoted at 15 μl and flash frozen in liquid nitrogen for preparation of grids for cryo-EM single particle reconstruction. Aliquots were kept frozen at -80 °C until grids were prepared.

### Cryo-EM grid preparation and data collection

The purified samples were applied to Quantifoil R 1.2/1.3 300 Mesh Gold grids. The grids were glow discharged for 90 s at 30 mA with a PELCO easiGlow™ Glow Discharge Cleaning System to obtain a hydrophilic surface. The glow-discharged grids were used to prepare vitrified samples with the Thermo Scientific Vitrobot Mark IV System. We applied 3 μl of purified protein to the glow-discharged surface of the grid at 4 °C at 100% of humidity and blotted the solution for 5.0 to 5.5 s, with blot force of either 18 or 19.

The data were acquired with a 300 kV Titan Krios G2 microscope (Thermo Fisher) equipped with a K3 Summit direct electron camera (Gatan) run in super-resolution mode at a nominal magnification of 105,000×, with a physical pixel of 0.834 Å. A phase plate was not used and the objective aperture was not inserted. SerialEM was used for automated data collection in beam-image shift mode with 9 images collected per stage movement, with a defocus range from -1.0 μm to -3.0 μm and beam-image shift compensation (15). The slit width of the GIF Quantum Energy Filter was set to 25 eV. Movies were dose-fractionated into 100 frames with a total dose of ∼80 e^-^/Å^2^. Two batches of data were collected. The first batch consisted of 2,398 movies, and the second batch consisted of 4,509 movies.

### Cryo-EM image processing and model building

All movies were imported into CryoSPARC followed by patch motion correction using binning of 2 and patch CTF correction (16). Particles were first picked using blob picker on 500 micrographs collected in the first batch to generate initial 2D templates. Templates were created from selected 2D classes and template picking on all micrographs from the first batch was conducted, resulting in a total of 1,498,114 particles. Particles were extracted with box size of 400 pixels followed by several rounds of 2D classification that yielded 77,806 clean particles. *Ab initio* reconstruction on the clean particle stack with three classes and one round of heterogeneous refinement was conducted and resulted in the monomer and the dimer initial classes, where the dimer class had a significantly lower resolution than the monomer one. Non-uniform refinement using a subset of particles for monomer and dimer resulted in a ∼4 Å resolution density map corresponding to the monomer structure and an unsuccessful dimer reconstruction (17). Topaz particle picker was used to further enrich particles that belong to each oligomerization state (18). Briefly, Topaz models were trained iteratively on a small set of particles with nearly unambiguous assignment (either all monomers or all dimers) from 25 micrographs. After each training, particles were extracted in 400-pixel boxes and cleaned by 2D classification. The newly cleaned particle stack re-entered the training until the model for monomer and dimer stopped improving. The models were then used to pick particles from all micrographs (each batch of data collected required separate Topaz training and extraction). Pooled monomer (779,882) and dimer (688,464) particles from two data collection batches were then processed separately via several rounds of 2D classification and heterogeneous refinement to remove impurities. A set of 150,028 monomer particles entered the final non-uniform refinement and resulted in a 3.15 Å resolution map; a set of 40,700 dimer particles entered the final non-uniform refinement with C2 symmetry and resulted in a map of 3.46 Å resolution. An initial model was obtained by docking the PDB 6WTI to the map using Molrep implemented in CCPEM (19-21). The model was then iteratively rebuilt manually in Coot and refined using *Phenix*.*real_space_refine* and Servalcat (22-29). The atomic coordinates (PDB: 8F68 and 8F6C) and maps (EMD-28877 and EMD-28879) have been deposited in the Protein Data Bank (http://wwpdb.org/) and the Electron Microscopy Data Bank (https://www.ebi.ac.uk/emdb/), respectively.

## Results

### 3D reconstruction of the ubiquinol oxidase monomer and dimer in the same sample

Initial 2D classification using particles picked from the same sample indicated the presence of both monomers and dimers of ubiquinol oxidases (**Figure 2A**). However, particle picking using blob or template-based methods favors the monomer, likely due to the higher abundance of monomers in the sample. To separately enrich particles for each oligomeric status, we used Topaz, a CNN-based particle picker (18). Several rounds of training and picking were performed, resulting in progressively purer sets of monomeric or dimeric particles at the end of each iteration, which were then used as the input for the next round of training. The final enriched sets of particles for each oligomeric states were then used in the next steps of the reconstruction process to obtain the final refined models. For the ubiquinol oxidase monomer, 150,028 particles were used in the final refinement and resulted in a 3.15 Å resolution map; for the dimer, 40,700 particles were used in the final refinement and the reconstructed map reached resolution of 3.46 Å. The data collection and image processing are summarized in **Table 1**. In both the monomer and dimer maps, the transmembrane domain of the protein was surrounded by an amphipol ring, consistent with lipid layer and/or nanodiscs positions in previously solved structures of this enzyme (PDB: 1FFT, 6WTI, 7CUW) (**Figure 2B and C**). Based on the number of particles in the final refined models, the monomer to dimer ratio in the sample is around 4:1, indicating a higher amount of dimer than the previous BN-PAGE observation (monomer:dimer = ∼50:1) (12).

**Table 1.**
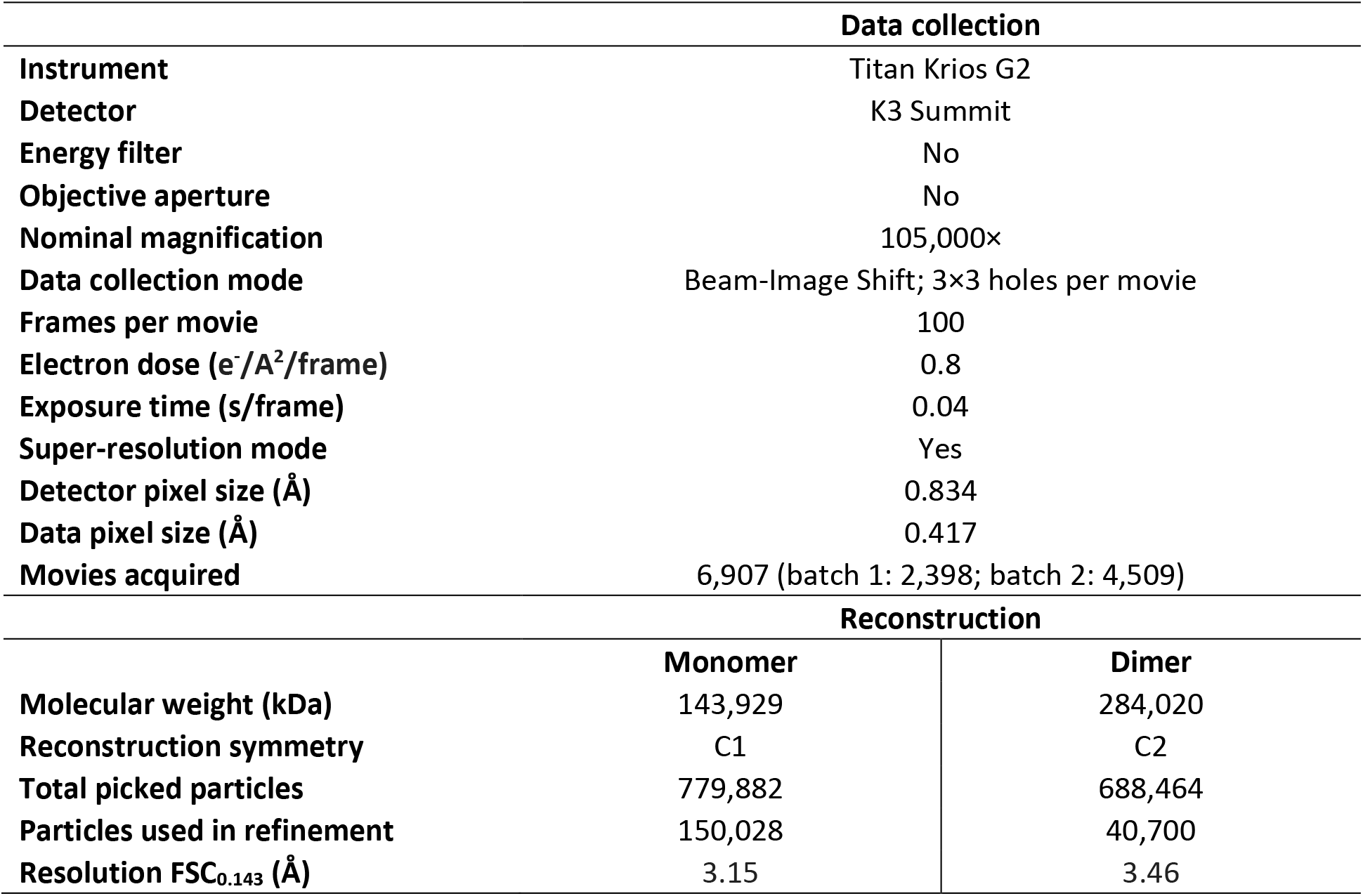
Data collection and processing.

**Figure 2.**
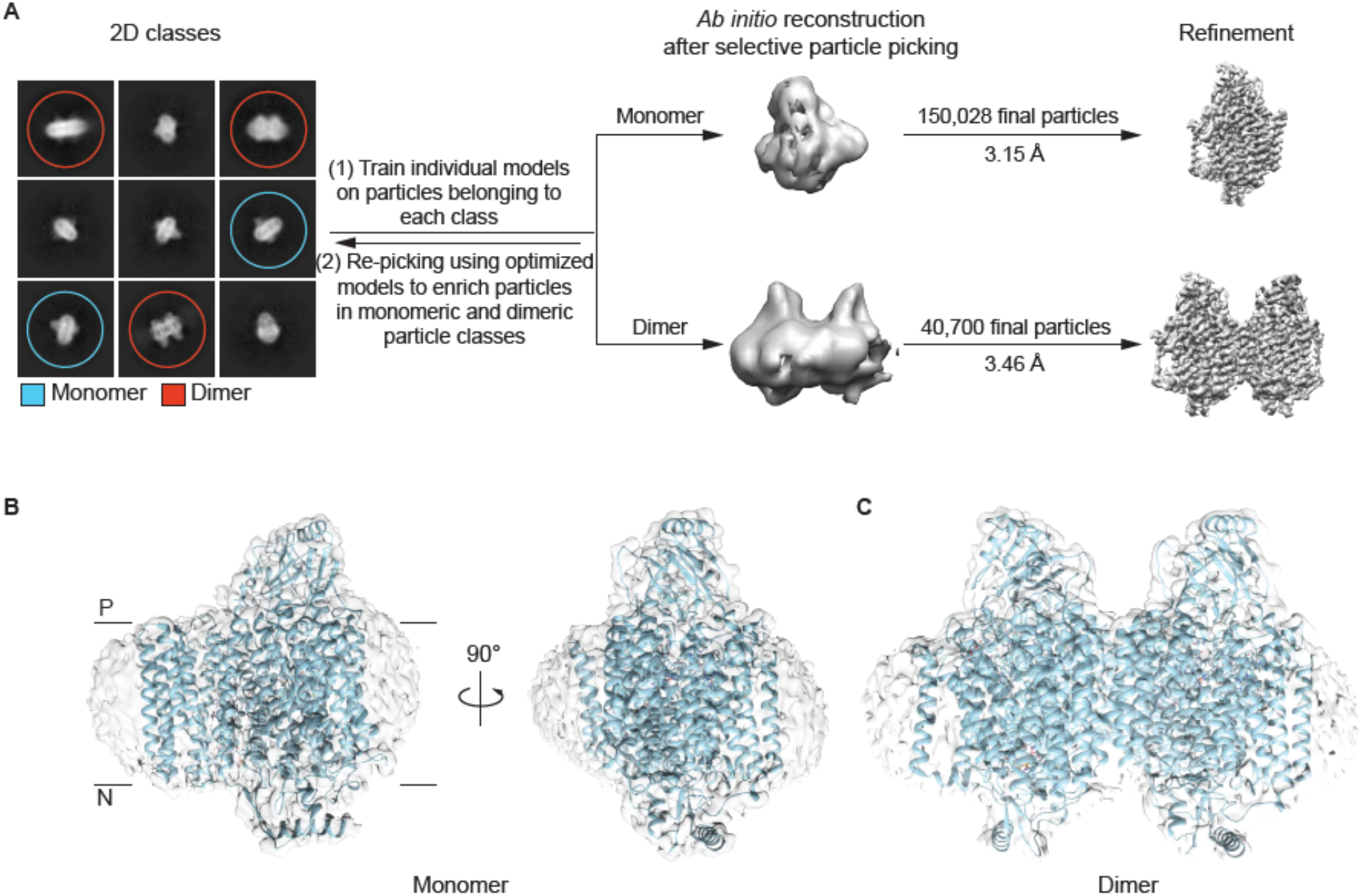
Cryo-EM reconstruction of the monomer and dimer structures of the *E. coli* ubiquinol oxidase. **A)** General workflow for cryo-EM reconstruction. Left panel: selected 2D classes represent both monomers (cyan circles) and dimers (red circles). Note that monomer and dimer 2D projections from certain angles may look similar to each other. Such classes are not labeled with a colored circle. Middle panel: *ab initio* reconstruction of the two ubiquinol oxidase classes. The dimer class has significantly lower abundance than the monomer class, so iterative AI-assisted particle picking was conducted to enrich the dimer class. Right panel: 3D maps of refined monomer (150,028 total particles, 3.15 Å resolution) and dimer (40,700 total particles, 3.46 Å resolution). **B)** Map and atomic model of the ubiquinol oxidase monomer. **C)** Map and atomic model of the ubiquinol oxidase dimer.

### Monomer structure of the ubiquinol oxidase

The structure of the ubiquinol oxidase monomer solved in this study is very similar to previously published results, with only differences in the number of bound ligands (**Figure 2B, Figure 3**). All four subunits, including residues 552-656 in subunit I which were missing in the original X-ray crystallographic structure (1FFT) showed clear density in the map of our reconstruction (EMD-28877) (**Figure 2B**) (1). Our atomic model (8F68) is comparable to the recently solved cryo-EM structures of this protein (7CUW and 6WTI) (2, 3), with the main chain RMSD between our model and either of the two other models (7CUW and 6WTI) around 0.4 Å (**Figure 3A**).

**Figure 3.**
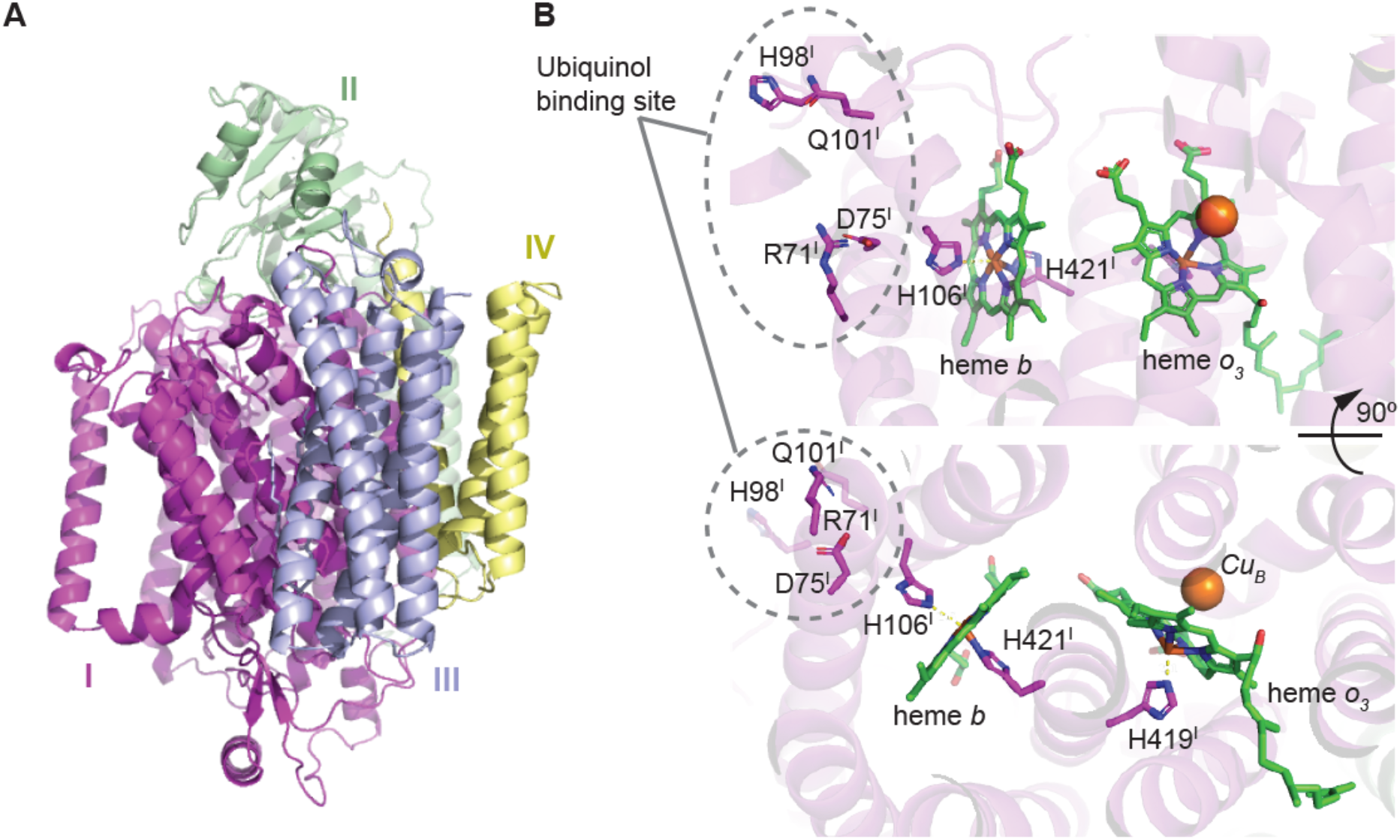
The *E. coli* ubiquinol oxidase monomer structure. **A)** Overview of the ubiquinol oxidase monomer structure solved in this study. The four subunits are labeled with color (I: magenta; II: green; III: blue; IV: yellow). **B)** Detailed view of the redox center from two different angles (bottom panel is top panel rotated 90 degrees). The ubiquinol binding site is labeled with a gray dashed circle and key residues are shown (R71^I^, H98^I^, Q101^I^). No ubiquinone molecule was observed in the structure. Heme *b* is coordinated with H106^I^ and H421^I^. Heme *o*_*3*_ is coordinated with H419^I^ and the copper ion is labeled as an orange sphere.

Both the heme *b* and the heme *o*_*3*_ molecules were observed in the map at expected locations. Specifically, the heme *b* molecule is coordinated by H106 and H421 from subunit I. Heme *o*_*3*_ is coordinated by H419 of subunit I on one side, and the copper ion on the other. However, there is no density consistent with ubiquinone at the ubiquinone binding site (**Figure 3B**), which leaves the redox center of the protein incomplete. In addition, the model in this study showed three 1,2-Distearoyl-sn-glycerophosphoethanolamine (3PE) molecules bound between subunit I and III, as observed in the published *E. coli* ubiquinol oxidase cryo-EM structures (2, 3). Other ligands observed in the previously solved cryo-EM structures, such as additional 3PE molecules (observed in 7CUW subunit I and in 6WTI subunit I, II and IV) and a pentadecyl(tetradecyl)peroxyanhydride (U9V) molecule (observed in 6WTI subunit I) were poorly resolved in our reconstruction, likely due to its lower resolution (3.15 Å), so they are not included in the atomic model. The 3PE and U9V ligands do not appear to be important for the function of the ubiquinol oxidase and will not be further discussed here.

### Dimer structure of the ubiquinol oxidase

The ubiquinol oxidase dimer (8F6C) has C2 symmetry, with the 2-fold axis oriented parallel to the transmembrane domains (**Figure 4A, C**). Local resolution analysis of the map (EMD-28879) shows that most regions of the reconstruction have resolution close to the global resolution of the map (3.46 Å), with only the map region corresponding to TM0 of subunit I (**Figure 4B**, in red dashes circle) having significantly lower resolution (> 4 Å), possibly due to higher flexibility arising from limited contacts between TM0 and the rest of the protein.

**Figure 4.**
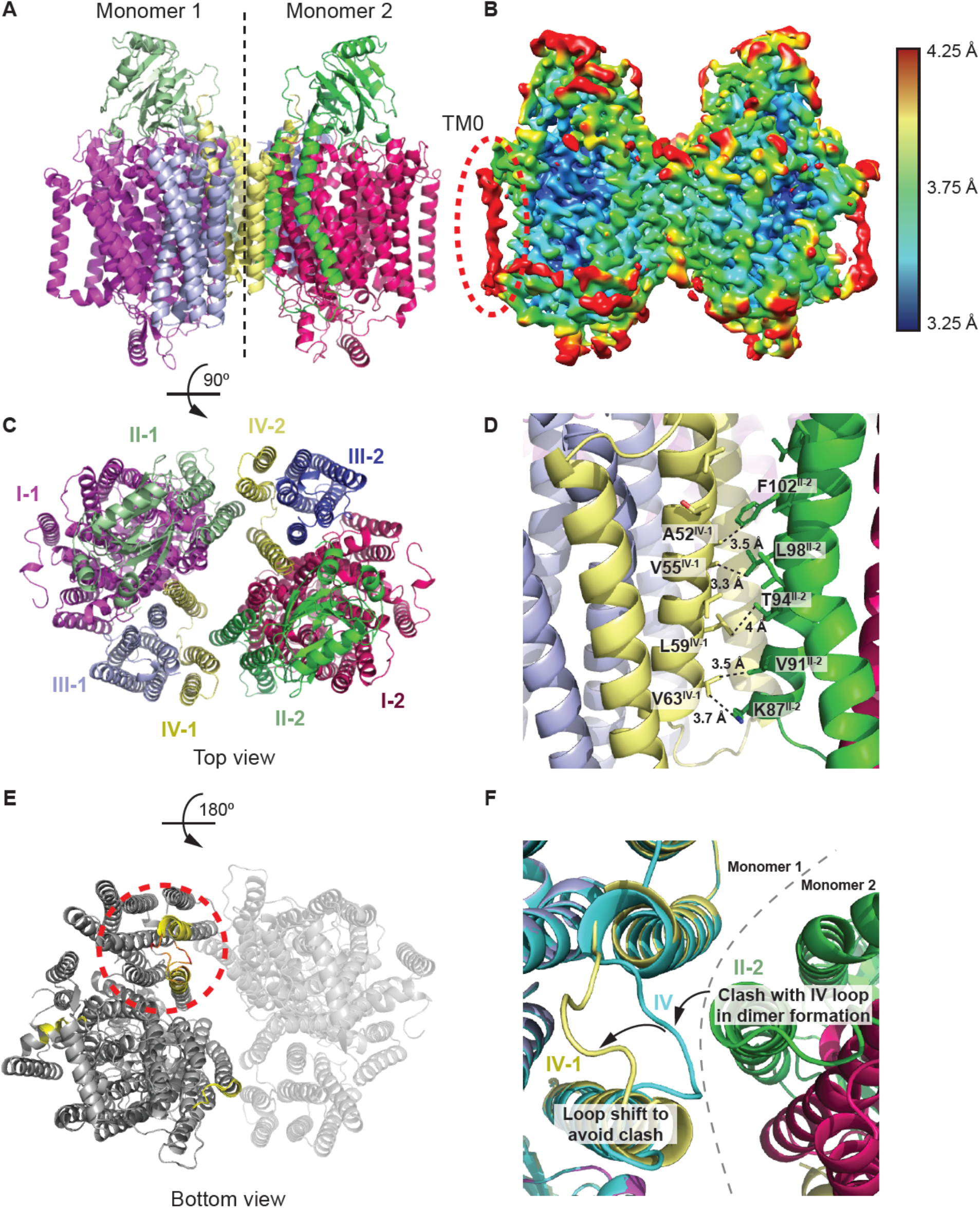
The *E. coli* ubiquinol oxidase dimer structure. **A)** Overview of the ubiquinol oxidase dimer structure solved in this study. The two monomers are separated by a dashed line in the middle. Color representation of the subunits is the same for monomer 1 as in Figure 3; brighter colors are used for monomer 2. **B)** Local resolution of the dimer map. TM0 in subunit I shows the lowest resolution among all regions. **C)** Top view of the dimer structure shown in panel A with each monomer and its subunits labeled in color. The dimer has C2 symmetry. **D)** Key residues at the dimerization interface. In the dimer, subunit IV of monomer 1 interacts through a series of hydrophobic interactions with the subunit II of monomer 2. Key residues involved in the hydrophobic interaction and the interaction distances are labeled. **E)** Bottom view of the dimer structure shown in panel A superimposed with the monomer structure solved in this study. Residues with superimposition RMSD lower than 0.3 Å are shown in grey, residues with superimposition RMSD higher than 0.3 Å are shown in a yellow-red gradient. Red dashed circle indicates the region of the protein with the highest RMSD between the monomer and the monomeric subunit of the dimer. **F)** Detailed view of the region with the highest RMSD between monomer and the monomeric subunit of the dimer. Subunit IV in the monomer structure is in cyan. Grey dashed line indicates the boundary between monomer 1 and monomer 2 in the dimer structure. Subunit IV of monomer 1 (IV-1) in the dimer structure is shown in yellow. Subunit II of monomer 2 (II-2) is shown in green. Arrows show the movement of the subunit IV loop position between the monomeric and dimeric structures. This movement is required so that clashes with subunit II of monomer 2 (II-2) can be avoided during dimer formation.

The dimerization interface is between subunit II of one monomer and subunit IV of the other monomer, with a large empty channel between the two monomers (**Figure 4C**). The total buried interface area between the two monomers is ∼556 A^2^, which is ∼0.6% of the total surface area of the dimer (87,166 A^2^) (30, 31). The dimerization interface is mostly non-polar, and the hydrophobic interactions occur along the helices of subunit IV of monomer 1 (yellow) and subunit II of monomer 2 (green), where the two helices are almost parallel to each other (**Figure 4D)**. Key residues that participate in hydrophobic interactions between the two subunits are shown in **Figure 4D**. Specifically, V63 from subunit IV, monomer 1 interacts with side chains of both K87 and V91 from subunit II, monomer 2. L59, V55 and A52 from subunit IV of monomer 1 form hydrophobic interactions with T97, L98 and F102 from subunit II of monomer 2. Beyond F102 of subunit II of monomer 2 (towards the C-terminus), the two parallel helices are too far from each other to engage in any interactions.

The monomeric subunit in the ubiquinol oxidase dimer structure (8F6C) is very similar to the ubiquinol oxidase monomer structure (8F68), with the main chain RMSD of 0.624 Å based on the alignment of 1204 residues (alignment on one monomer). **Figure 4E** shows the superimposition of the monomer and the dimer structure; regions with superimposed RMSD higher than 0.3 Å are highlighted in a gradient of yellow and red. The most significant difference is located in the loop in subunit IV (residues 67–74) (**Figure 4E**, red dashed circle, **Figure 4F**). Before the dimer formation, this loop swings outward to the surface of the protein (**Figure 4F**, cyan). After the dimer formation, subunit II of monomer 2 forces this loop to move away from the dimer interface to avoid a clash (**Figure 4F**, yellow). The maximum shift of this loop is ∼12 Å between the monomer and dimer state.

### Discussion and conclusion

In this study, we report two *E. coli* cytochrome *bo*_*3*_ ubiquinol oxidase structures. The monomer structure was reconstructed at 3.15 Å, a lower resolution than the published cryo-EM structures 7CUW (resolution 2.63 Å) and 6WTI (resolution 2.38 Å) (2, 3). The dimer structure was reconstructed at an intermediate resolution of 3.46 Å and is the first dimer structure reported for this protein.

A few points are worth mentioning about the structures solved in this study in comparison to the published structures (1-3). Firstly, amphipol was used in this study to maintain the solubility of the membrane protein, instead of detergent, styrene maleic acid co-polymer (SMA) or nanodiscs. The use of amphipol does not seem to affect the overall structure of the monomer, as supported by the low RMSD when superimposed with the previously reported structures. Secondly, both the monomer and the dimer structures lack the ubiquinone molecule bound to subunit I, which results in the incomplete redox center. It has been reported that certain detergents such as Triton X-100 could strip the bound ubiquinone from the protein (10). Only DDM and amphipol, which are routinely used in purification of similar ubiquinone-bound enzymes were used during purification in our study, so it is unclear at which step ubiquinone was lost from the protein (2, 32). Although the ubiquinone was missing in the structures, the lack of this cofactor is unlikely to be the reason for the dimerization, because: 1) the ubiquinone binding site is relatively distant from the dimerization interface; and 2) the ubiquinone is missing in the monomeric X-ray structure of this enzyme (1FFT) (1). DDM was also used in the study by Stenberg, *et al*. where the ubiquinol oxidase dimer was observed on BN-PAGE, suggesting that DDM may preserve the dimer better than other detergents (e.g., Triton X-100) (11, 12). Thirdly, the measured dimerization interface is relatively small and covers only a small portion of the entire protein surface. However, this does not necessarily mean that the dimer structure is an experimental artifact. Such small dimerization interfaces have been observed in other complexes/structures solved by cryo-EM (33). Further experiments are required to determine whether the dimer is functional and/or physiologically relevant.

To summarize, in this study we solved the structures of the *E. coli* ubiquinol oxidase in two oligomerization states (monomer and dimer) from the same sample. The monomer structure, despite its intermediate resolution and the lack of ubiquinone cofactor, is highly consistent with the other published results and adds support to previously observed orientation for residues 552-656 in subunit I that was missing in the X-ray crystallographic structure (1FFT) (1). The dimer structure shows that the protein dimerizes in C2 symmetry through mainly hydrophobic interactions between subunit II from one monomer and subunit IV from the other monomer, with minor structural changes in the other parts of the protein. Since it is the first dimer structure solved for the *E. coli* ubiquinol oxidase, it provides the groundwork for future hypothesis testing (e.g., sites for mutagenesis) regarding the physiologically relevant oligomerization status and functions of this protein.

## Acknowledgements

This project has been funded in whole or in part with federal funds from the National Institute of Allergy and Infectious Diseases, National Institutes of Health, Department of Health and Human Services, under Contracts HHSN272201700060C and 75N93022C00035. This project was also supported by funds from the Department of Energy (DE-SC0019600 to YG and ZO, DE-SC0021600 to ZO) and funds from the National Institute of General Medical Sciences, National Institutes of Health (R35GM145365 to ZO).

We thank the Cryo-Electron Microscopy Facility (CEMF) at UT Southwestern Medical Center which has been supported by grants RP170644 and RP220582 from the Cancer Prevention & Research Institute of Texas (CPRIT) for maintaining a Titan Krios microscope.

## Author Contributions

YG, EK, ZO, DB and AS designed the research. YG, EK, TE and DB generated data; YG, EK, ZO, DB and AS analyzed data; YG, EK, DB and AS wrote the paper.

